# Optimal network sizes for most robust Turing patterns

**DOI:** 10.1101/2024.10.15.618426

**Authors:** Hazlam S. Ahmad Shaberi, Aibek Kappassov, Antonio Matas-Gil, Robert G. Endres

## Abstract

Many cellular patterns exhibit a reaction-diffusion component, suggesting that Turing instability may contribute to pattern formation. However, biological gene-regulatory pathways are more complex than simple Turing activator-inhibitor models and generally do not require fine-tuning of parameters as dictated by the Turing conditions. To address these issues, we employ random matrix theory to analyze the Jacobian matrices of larger networks with robust statistical properties. Our analysis reveals that Turing patterns are more likely to occur by chance than previously thought and that the most robust Turing networks have an optimal size, surprisingly consisting only of a handful of molecular species, thus significantly increasing their identifiability in biological systems. This optimal size emerges from a tradeoff between the highest stability in small networks and the greatest instability with diffusion in large networks. Furthermore, we find that with multiple immobile nodes, differential diffusion ceases to be important for Turing patterns. Our findings may inform future synthetic biology approaches and provide insights into bridging the gap to complex developmental pathways.

## Introduction

Spatial patterns and structures are ubiquitous in biological systems, ranging from microbial communities and biofilms to developmental biology and ecological systems [1] [2] [3]. In seminal work, Alan Turing (and later Gierer-Meinhardt) proposed a self-organizing emergent mechanism for pattern formation based on a diffusion-driven instability, utilizing a slowly diffusing activator and a fast-diffusing inhibitor molecule [4] [5]. According to Turing’s definition, the chemical reactions are stable without diffusion, leading to a homogeneous steady state, but become unstable with diffusion, forming a periodic pattern at certain wave numbers. The mathematical analysis is generally based on reaction-diffusion models, implemented using partial differential equations, and linear stability analysis to investigate the effect of perturbations on a steady state [6]. Nonetheless, employing Turing models to understand biological patterns is still a topic of debate.

The Turing mechanism clearly has biological relevance in cell and developmental biology, likely explaining aspects of digit formation in mice, zebrafish skin, fingerprint formation, and cortical folds of the fetal brain [7][8][9][10]. However, this mechanism has two main drawbacks: extreme simplicity and a need for fine-tuning. In contrast to simple activator-inhibitor Turing models, biological gene regulatory networks in embryonic development generally consist of hundreds of molecular species and are notable for their immense complexity, hierarchical structure, and tolerance to noise [11]. In contrast, the Turing conditions lead to a lack of structural robustness, which is at odds with the noise tolerance and evolutionary adaptability required for such patterning solutions to occur [12][13][14]. However, there are also bottom-up approaches to further elucidate the issues with Turing models.

As developmental biology is complex, synthetic biology provides an alternative route, following Feynman’s mantra: “What I cannot create, I do not understand” [15]. By implementing the Turing mechanism from scratch in cells that communicate via chemicals, our understanding of how to generate stable pattern can be systematically improved. Two previous synthetic biology attempts failed due to the lack of differential diffusion, leading to highly irregular patterns. First, stochastic Turing patterns were engineered in *E. coli* cells. The implemented circuit contained self-activation and lateral inhibition, with two diffusible quorum-sensing signals [16]. Second, solitary patterns were engineered in HEK cells using the Nodal–Lefty system [17]. The issue is that small molecules have roughly the same diffusion constant, although extra nonspecific binding can help slow down diffusion. In recent work, a more robust three-node network was implemented using synthetic circuits of six genes with small diffusible quorum-sensing molecules and extra control cassettes [18]. This showed regular patterns in growing bacterial colonies, but the variability is immense, and robustness is still limited. Hence, while there is progress in the rational design of circuits with specific properties, the lack of control over robustness represents a significant downside in tissue engineering, patterned biomaterial deposition, and bridging developmental programs [19][20][21].

The need to develop theoretical frameworks beyond the two-equation Turing model has been recognized previously. An exhaustive exploration of all two- and three-node topologies showed that three nodes are, on average, more robust than two nodes, pointing toward complexity increasing robustness [22]. More specifically, approximately 60% of all topologies produced patterns for some parameter combinations, but the parameter space producing Turing patterns was overall minuscule, around 0.1%. This finding is supported by another study that explored networks with up to four nodes [23]. Motivated by Robert May’s work in theoretical ecology [24] (based on earlier work by Eugene Wigner [25]) a random matrix approach for *N* ≤ 6 showed that larger networks add robustness by reducing the “diffusive threshold”, i.e., softening the requirement of differential diffusion [26]. This raises the question: why not investigate even larger matrices? While the diagonalization of Jacobian matrices is a slow *O*(*N* ^3^) process, it is still efficient for significantly larger networks with modern computers. The advantages of exploring random Jacobian matrices are apparent; such an approach produces excellent statistics, and biological realism can be introduced through the distributions and topologies.

Here, we go beyond small Turing networks and use large random Jacobian matrices to systematically explore how network size affects robustness. We begin by motivating distributions for Jacobian matrix elements from explicit small Hill-function-based models for gene regulation, as relevant to synthetic biology [18]. We then continue by exploring large random networks with up to *N* = 100 nodes using corresponding random Jacobian matrices with two diffusers and variable sparsity. We identify an optimal network size *N*_opt_ ∼ 5 − 8, arising from a tradeoff between stability without diffusion and instability with diffusion, each with opposite *N* -dependencies. This increased robustness relieves the constraints on parameters, known as Turing conditions, including differential diffusion for the two diffusing species. Our results are expected to renew the quest for robust Turing patterns in synthetic biology and to identify Turing modules in larger networks of developmental biology.

## Results

We are interested in the robustness of large Turing networks, that is reaction-diffusion models with many molecular species with a large parameter space to support diffusion-driven instabilities. Fig. 1 shows example networks, ranging from small (*N* = 2 nodes) to large (*N >>* 1). We restrict ourselves to two diffusing species similar to Turing’s original paper. In order to understand how to model large networks we begin by investigating examples of smaller networks to gain intuition. For a network of *N* nodes and hence *N* species with time and space dependent concentrations ***X***(***r***, *t*) = { *x*_1_(***r***, *t*), *x*_2_(***r***, *t*), …, *x*_*N*_ (***r***, *t*)}, the general dynamics are given by the partial differential equations

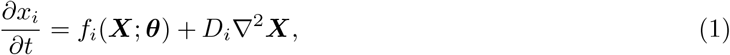

where ***D*** = *diag*(*D*_1_, *D*_2_, 0, …, 0) is the diagonal diffusion matrix, and Laplacian 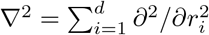 in *d* dimensions, given by the sum of second spatial derivatives. For simplicity, we assume an infinite domain, allowing us to neglect boundary effects and to use a continuous wave number. As our motivation is to learn how to build these with synthetic gene circuits, we model the activating and inhibiting interactions with Hill functions, along with basal expression and degradation:

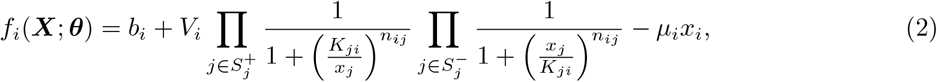

with ***θ*** = {***b, V***, ***K, n, µ***} respectively the sets of basal and maximal expression rates, concentration thresholds, Hill coefficients, and degradation rates. Furthermore, 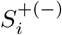 denotes the set of positive (negative) edges ending in node *i*, where the different regulators act (and saturate) independently (non-competitively) of each other. Following [22], we sample parameters from a wide range of allowed values in arbitrary units.

**Figure 1:**
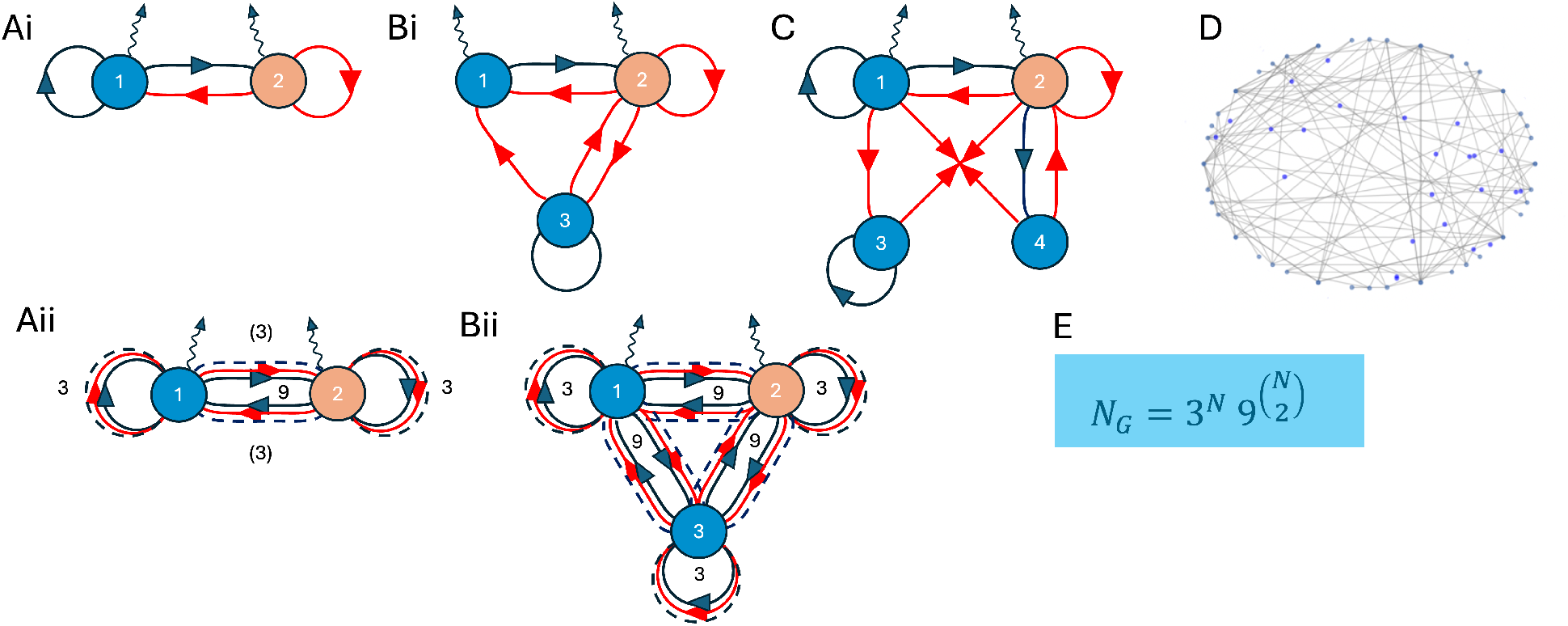
Network graph representations of pre-defined Turing topologies. (Ai) 2-node Gierer-Meinhardt network. (Bi) 3-node network based on [22, 18]. (C) Extended 4-node network. (D) Large *N >>* 1 network. (E) Number of different Turing topologies (graphs) for *N* nodes, exemplified by all 2 and 3-node networks in (Aii) and (Bii), respectively. Dark blue (red) arrows indicate activation (inhibition), and wiggly lines indicate diffusion.

To understand whether such parameter combinations produce Turing patterns, we use linear stability analysis following Turing’s approach (see Methods for additional details). Briefly, first the model needs to have a stable homogeneous steady state ***X***^*\**^ without diffusion, defined by

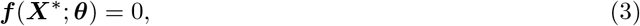

which we solve with the Newton-Raphson method using different initial conditions. Second, we linearize the dynamic equations assuming small perturbations around steady state, using 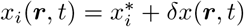, leading to

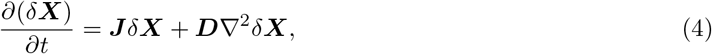

with Jacobian matrix ***J*** of first derivative with matrix elements *J*_*ij*_ = *∂f*_*i*_*/∂x*_*j*_.

To get rid off the second derivatives, we Fourier transform in the space domain, or more simply apply a wave-like perturbation 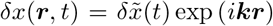 exp (*i****kr***) with ***k*** = {*k*_1_, *k*_2_, …, *k*_*d*_) and ***kr*** the scalar product between the ***k*** and ***r*** vectors. Plugged into Eq. 4, this leads to a new Jacobian matrix with modified diagonal matrix elements and an extra parameter *k* = |***k***|

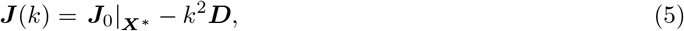

indicating that for a rotationally invariant system, the dependence is only on the modulus of ***k***, rendering linear stability analysis an effective one-dimensional problem. As a result, diagonalization of the modified Jacobian matrix leads to a *k*-dependent eigenvalues, called dispersion relations, where the one with the largest real part, termed *Re*(*λ*_max_(*k*)), determines the stability. For a Turing instability, we require the homogeneous steady state without diffusion to be stable (corresponding to *k*=0), i.e. with a negative real part given by *Re*(*λ*_max_(*k*(0))) *<* 0. Furthermore, with diffusion we require instability for a wave number (*k >* 0) and thus a positive real part given by *Re*(*λ*_max_(*k*)) *>* 0. For a classic Turing instability leading to a well-defined wave pattern, there is a *k*_max_ *>* 0, for which the dispersion has a clear maximum, and for *k* → ∞, the system becomes stable again (negative real part of eigenvalue). This type of instability is called Turing I (Fig. 2A). However, there are other possibilities, such as instability for *k* → ∞ (Turing II in Fig. 2B). (Note, this is a simplified classification scheme - in other schemes our Turing I is given by Turing Ia, and our Turing II is given by Turing Ib, IIa, and IIb with subtle differences [22].) Yet, another type of instability is the Turing-Hopf, for which the imaginary part can additionally lead to oscillations (not shown).

**Figure 2:**
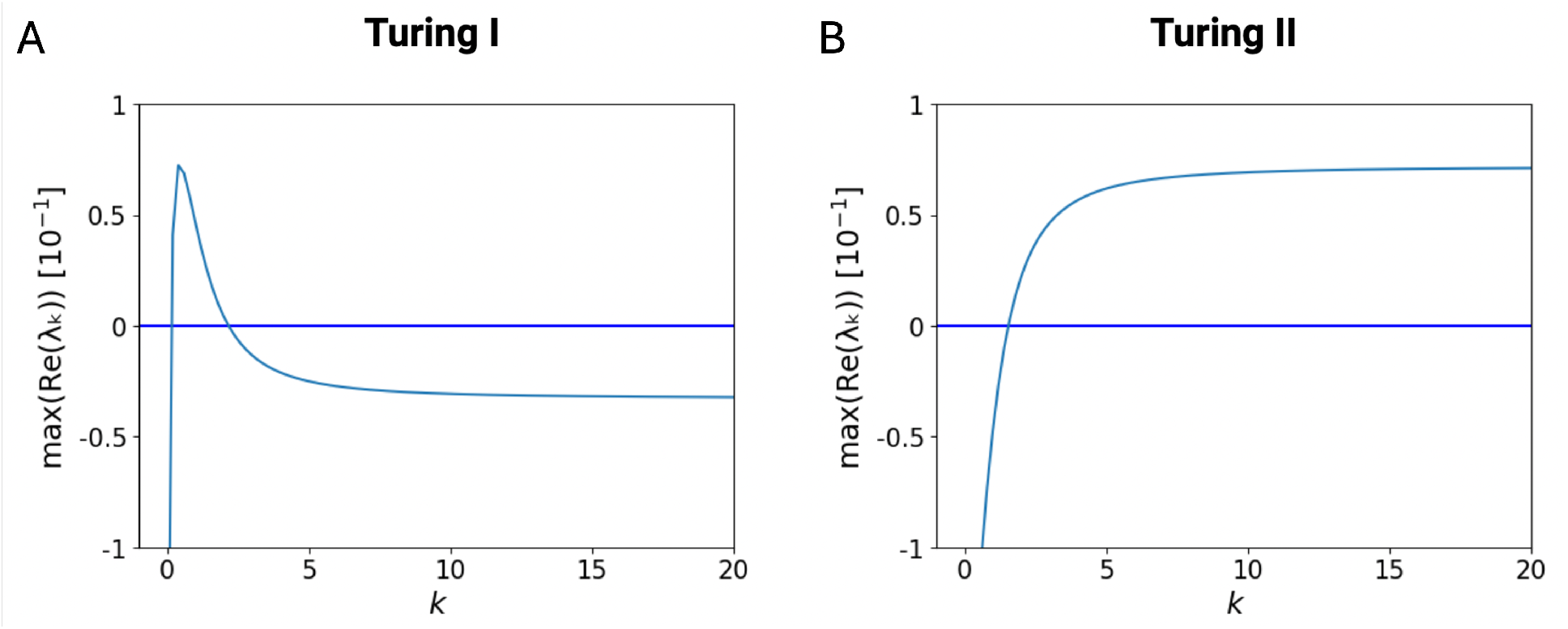
Dispersion relations illustrating different Turing instabilities. (A) Turing type I instability with a peak and negative real part for *k >>* 1. (B) Turing type II instability which stays positive (note this is however not possible when all species diffuse). The *y* axis represents the maximum real part of the eigenvalue and the *x* axis shows wave number *k*. The plots are generated from random matrices with network size *N* = 10 and variance *σ*^2^ = 0.25.

Focusing on small networks (*N* = 2 − 4), we used Latin hypercube sampling of 10^7^ parameter sets and filtered these for Turing I instabilities. Fig. 3 shows the fits of the empirical histograms of the Jacobian matrix elements (*j*) to the beta distribution *B*(*j*; *α, β*) (see Methods for details on parameter searches and fitting procedure). The beta distribution depends on two parameters, *α* and *β*, allowing a wide range of distributions from symmetric Gaussian-like to non-Gaussian shapes with various levels of skewness and kurtosis. We generally observe that the Turing-I matrix elements are more narrowly distributed in line with Turing I having to fulfill Turing conditions, leading to a subset of parameters and hence Jacobian matrix elements. This can clearly be seen in Fig. 3A (see Fig. S1 in Supplementary Materials for additional histograms). To fulfill the Turing conditions, the (1,1) Jacobian matrix elements need to be positive (activator), which is the case for the histogram and fitted line (blue), but not for the orange line, representating the non-Turing case. Furthermore, the (2,2) matrix elements need to be negative (inhibitor), which is here already fulfilled for non-Turing due to model constraints (degradation). In contrast, the (1,2) and (2,1) matrix elements can respectively be either negative and positive (here the case even for non-Turing due to model constraints), or the other way around (however not for our 2-equation model) [6]. We generally did not observe multimodal distributions, pointing to a certain level of simplicity in the distributions.

**Figure 3:**
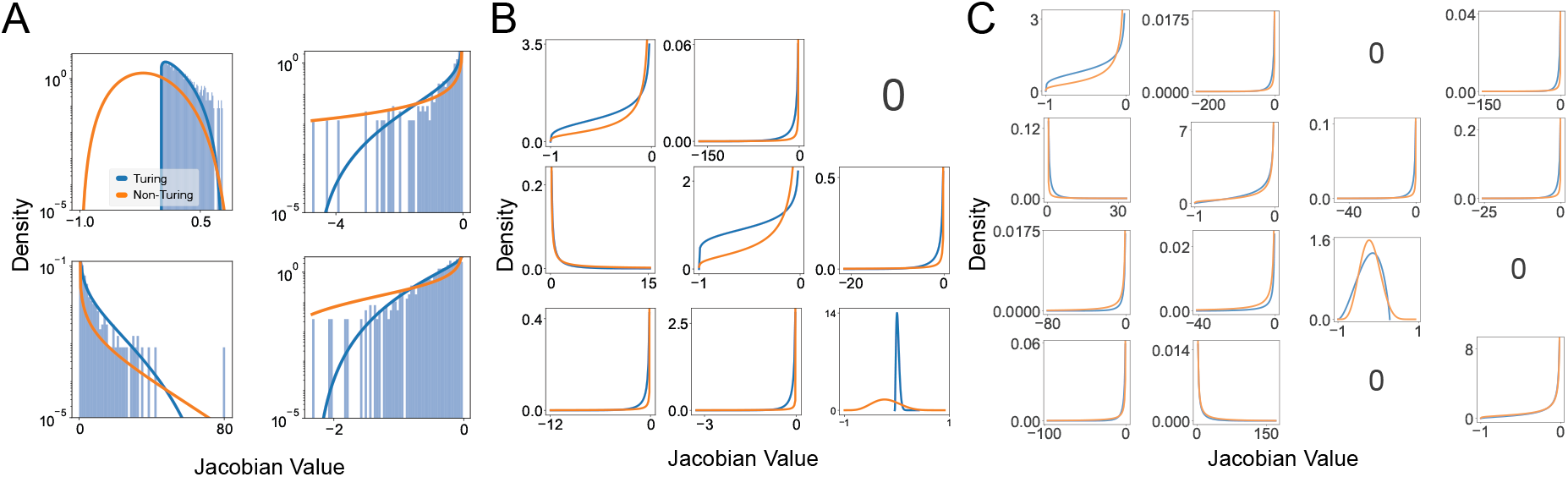
Empirical distributions of Jacobian matrix elements. (A) Histograms for parameters leading to Turing I instabilities (light blue) and fits to the beta distribution for Turing I (blue line) and non-Turing (orange line). Empirical distributions are computed based on parameter sampling for the pre-defined 2-4-node network topologies in Fig. 1Ai, Bi, and C for 2-node (A), 3-node (B), and 4-node (C) networks. Note some matrix elements are zero due to missing links in the network (or entries in the adjacency matrix).

The difference between Turing and non-Turing Jacobian matrix elements is further visible in Fig. 4, where panel A shows beehive plots based on the empirical data from Fig. 3. Clearly, the variances of the off-diagonal matrix elements of the non-Turing cases are much broader than the Turing-I cases. This is further provided in panel B, showing the sum of the Jacobian matrix elements (equivalent plots can be made for the mean of those random variables). For larger matrices, we expect to observe the central limit theorem, with the sums calculated from *N* (*N* − 1) off-diagonal Jacobian matrix elements. For increasing matrix size, and this is already evident for *N* = 4, we obtain approximate Gaussian distributions (still with a sharp peak at zero): The mean is approximately zero, the variance approaches a finite value (as variances are positive numbers), and the skewness and kurtosis vanish. Hence not surprisingly, on average a Gaussian distribution with zero mean and finite variance describes the Jacobian matrix elements. Note this trend would be even stronger, if we average over all possible network topologies with *N* nodes, which scales as 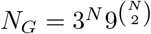 and hence is super-exponentially increasing with *N* (see Fig. 1E). In the following we exploit these insights and use random Jacobian matrices directly.

**Figure 4:**
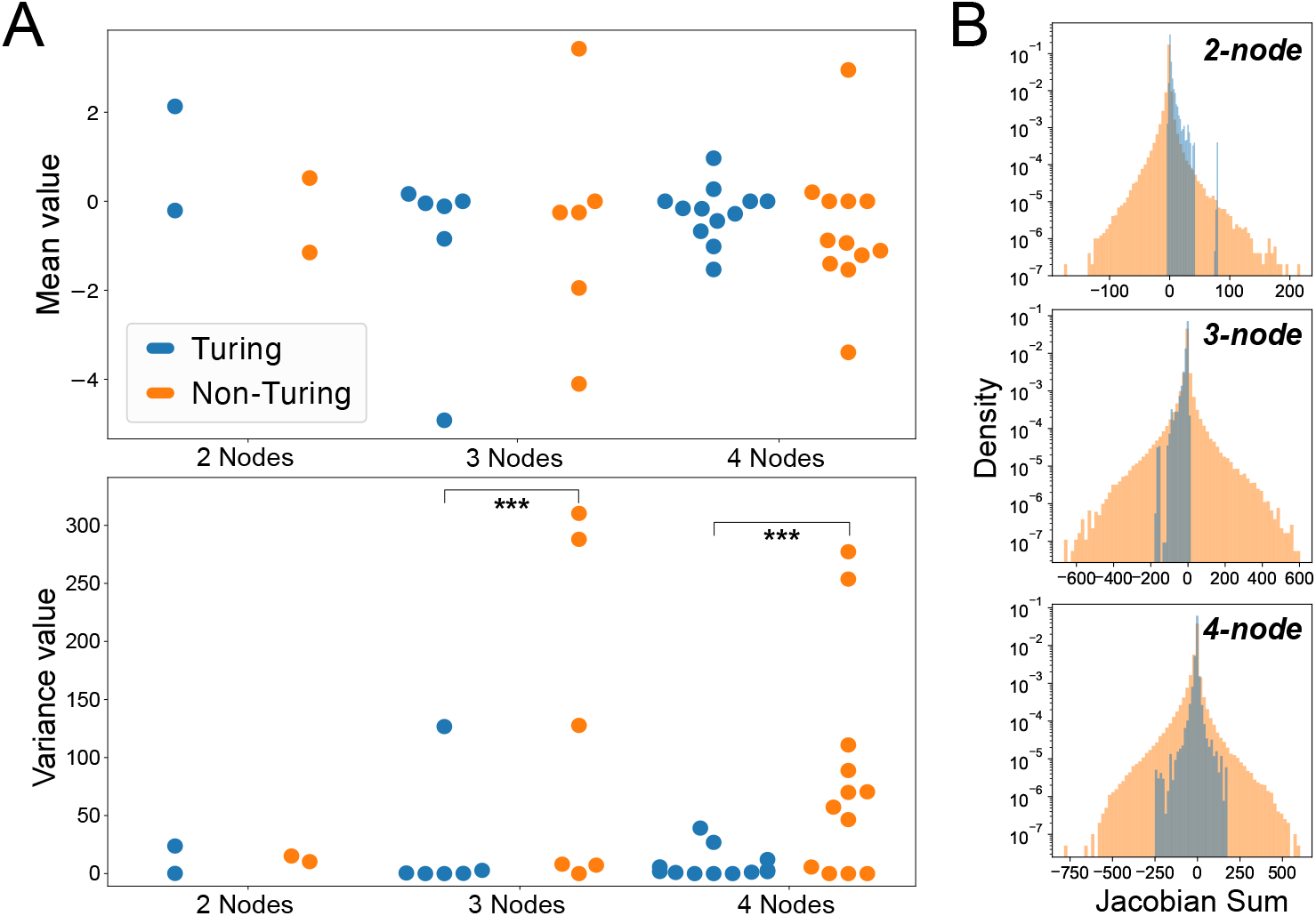
Jacobian matrix elements are different for Turing I and non-Turing. (A) Beehive plots of the mean (top) and variance (bottom) of the empirical distributions of the non-zero off-diagonal Jacobian matrix elements from Fig. 3 (based on the pre-defined 2-4-node network topologies in Fig. 1Ai, Bi, and C). A statistical Brown-Forsythe test was utilised to verify whether the difference in variance between each pair of Turing and non-Turing distributions was statistically significant. For all pairs of the Jacobian distributions except one, we found that the test showed a statistically significant difference between the variances. Furthermore, 12 out of 16 *p*-values were under 0.001, showing a strong statistical significance. (B) Sum of off-diagnal matrix elements, showing a clear visual difference between Turing I (light blue) and non-Turing (light orange) for the 2-4 node topologies. The distributions become increasingly symmetric (and Gaussian) for more nodes.

Inspired by Robert May’s statistical treatment of non-equilibrium ecological communities that studied asymmetric random Jacobians to represent linearized population dynamics, as opposed to Wigner’s symmetric (Hermitian) random matrices [24][27], we analyzed Turing instabilities by sampling asymmetric random Jacobian matrices that represent the linearized reaction dynamics. Similar to May’s “neutral interaction” model, our Jacobian matrix without diffusion is given by

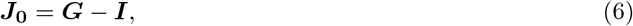

where ***G*** is a random matrix with elements *g*_*ij*_ randomly assigned from a Gaussian distribution that has mean value zero and variance Var(*g*_*ij*_) = *σ*^2^, but with diagonal terms *g*_*ii*_ = 0. For simplicity (and convenience), the latter are modeled by the identity matrix, ***I***, responsible for providing stability. In such a Turing system, a positive value of *J*_*ij*_ (or *g*_*ii*_) implies activation while a negative value implies inhibition, similarly to previous matrix representations of Turing systems [22][28]. The identity matrix ***I*** represents the degradation of the species. Further self-interactions are neglected, leading to the curious result that there are no Turing instabilities for *N* = 2 (see proof in Methods). While the Gaussian distribution is motivated by Fig. 4, there is no reason to expect the reaction kinetics in a reaction-diffusion system to be independent as there is no reason for the population dynamics in the work of May [24] to be independent (or normally distributed). Nevertheless, in the absence of experimental understanding of what the distribution of the parameters should be, the potential of the random matrix approach to explaining stability principles is greatly evidenced for population dynamics [29, 30] and in small Turing systems (*N ≤* 6) [26].

Next, we investigated the eigenvalue distributions of such random matrices (see Fig. 5A). In the absence of diffusion (i.e. at *k* = 0), the eigenvalues are distributed in a circle with radius 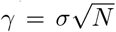 according to May’s “circular law”, derived from random ecological communities [24][27]. In the presence of diffusion, the eigenvalue distribution can be described as having “in-bulk” and “outlier” distributions (see Fig. 5B). The outlier distribution has extreme negative real parts, following approximately 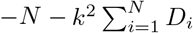 (see Methods), which tends to minus infinity for increasing *k* and network size *N*. As we are only interested in the maximal real part to determine the linear stability, it is safe to ignore the “outliers” and only focus on the “in-bulk” eigenvalue distribution where the maximal real eigenvalues are located in.

**Figure 5:**
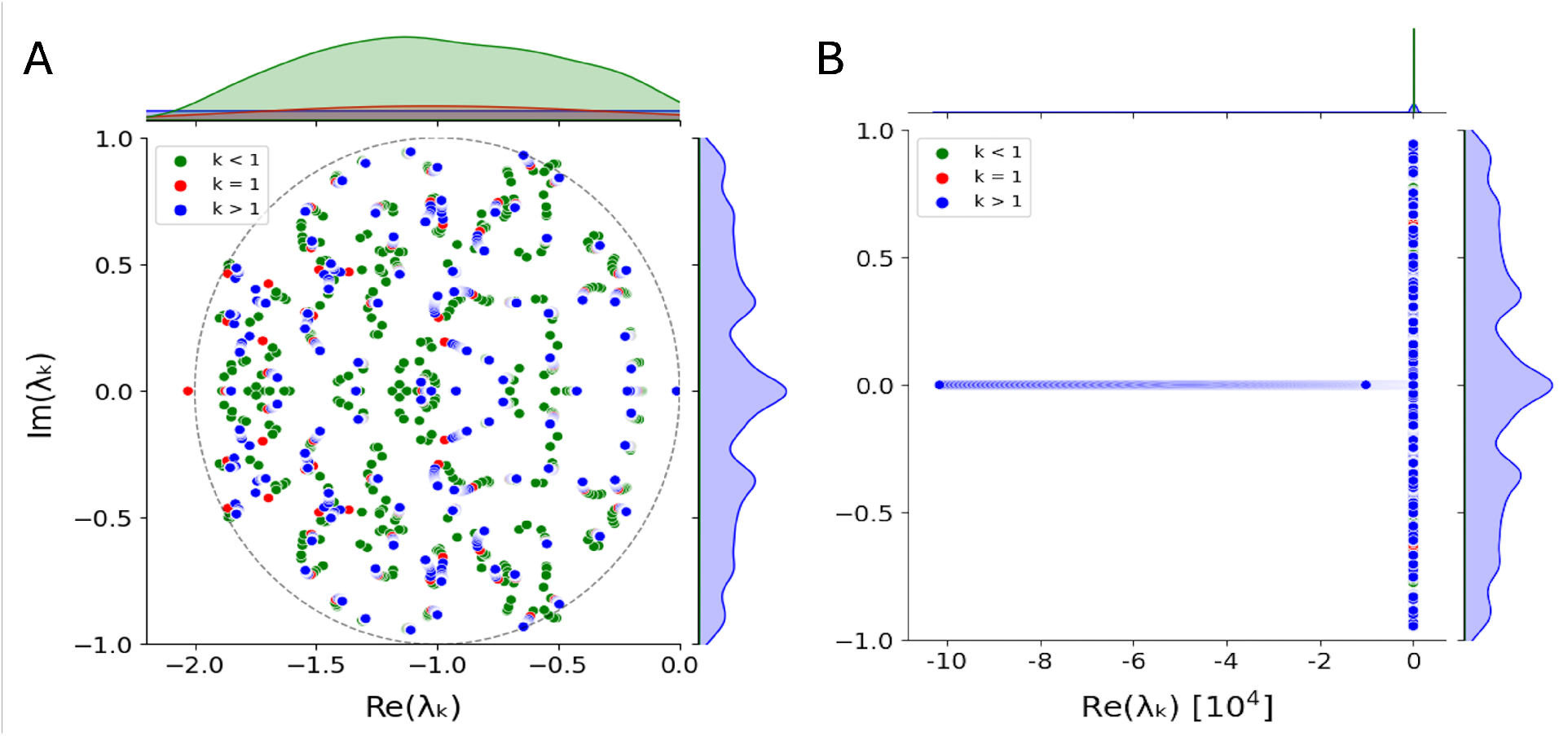
Eigenvalue distribution of a large random network. (A) Circular scatter plot and corresponding kernel density estimations of eigenvalues in the complex plain for network size *N* = 100, variance *σ*^2^ = 0.01, and diffusion constants *D*_1_ = 1, *D*_2_ = 10. While the system without diffusion has all the eigenvalues distribute around *γ* = 1, the same system with diffusion has most but not all of its eigenvalues distribute in a circle with radius *γ* = 1. (B) Full eigenvalue distribution with outliers. Green, red and blue points correspond to wave numbers *k <* 1, *k* = 1 and *k >* 1, respectively.

We subsequently sampled thousands of random matrices to analyse how network size affects the stability of a reaction-diffusion system. We observed three relationships between network size and stability. Firstly, in the absence of diffusion, a random reaction-diffusion system with fixed radius *γ* is likely more stable for smaller network size (Fig. 6A). Asymptotically, the circlar law predicts a step function for *N* → ∞, i.e. stability for radii *γ <* 1 (as circles are fully to the left of the imaginary axis) and instability for *γ >* 1 (as circles crosses the imaginary axis) [25]. Secondly, a previously stable system without diffusion is more likely to turn unstable (i.e. have a diffusion-driven instability) when diffusion is present for larger network size (Fig. 6B). Therefore, thirdly, out of all random matrices, Turing instabilities are maximized (i.e. found the most) at an optimal intermediate network size that is not too small or not too large (Fig. 6C).

**Figure 6:**
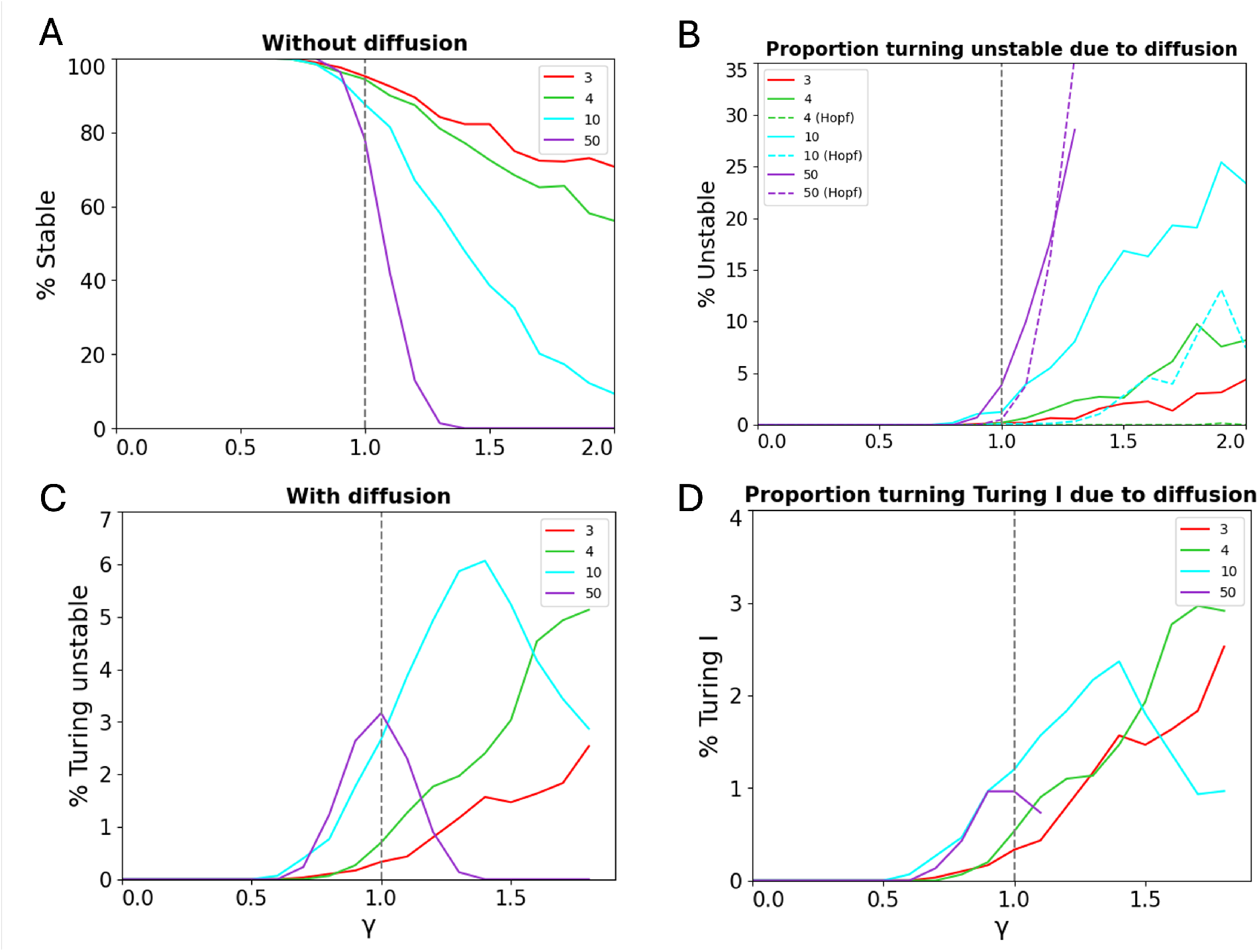
Relationships between network size *N* and radius 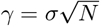. 10^3^ random matrices are sampled for each *N* and *γ* pair. (A) Percentage of stable random matrices without diffusion. (B) Percentage of previously stable random matrices turning unstable with diffusion. ‘Hopf’ indicates Turing-Hopf. (C) Percentage of random matrices leading to Turing instabilities (Turing I, II, and Turing-Hopf). (D) Percentage of random matrices leading to Turing I instabilities.

As the Turing-Hopf instability is competing with the Turing instability, and Turing I is of importance in producing spatial patterns, we analysed the effects of network size on Turing I and Turing-Hopf instabilities. Out of stable systems without diffusion, the systems are likely to become Turing-Hopf unstable in the presence of diffusion with larger network size (Fig. 6B). This trend for the Turing-Hopf instability is similar to the trend for Turing I. However, particularly at a very large network size, the Turing-Hopf instability becomes more likely to occur and ultimately out-competes Turing I, as can be seen for *N* = 50 in the figure. A similar trend is observed for Turing I relative to the overall Turing instabilities of all types: Turing I is more likely for larger network size being approximately half the occurrence of all Turing instabilities. Among all Turing instability types, we found that Turing II is the most common to arise (4.2% maximum occurrence of all random reaction-diffusion systems), followed by Turing I (3.6%).

Without having an experimental understanding of the variability of interaction strength between nodes, an optimal network size should produce Turing I instabilities for a wide range of variances. Such a network size would be most robust to changes in interaction strength, which is a desirable outcome from a biological perspective. To find the exact network size that is the most optimal, we plotted the occurrence of Turing instabilities for different network sizes and variances from *N* = 3 to 100 (Fig. 7A). Based on the heat map, the majority of the Turing instabilities arose just above *γ* = 1, as we should expect since *γ >>* 1 would likely result in an unstable system without diffusion, but *γ <<* 1 would make it harder for a stable system to become unstable with diffusion. In fact, the occurrence of Turing is “pinned” to radius *γ* = 1 for large *N*, and the percentage density then follows closely the circular law *σ*^2^ ∼ 1*/N*.

**Figure 7:**
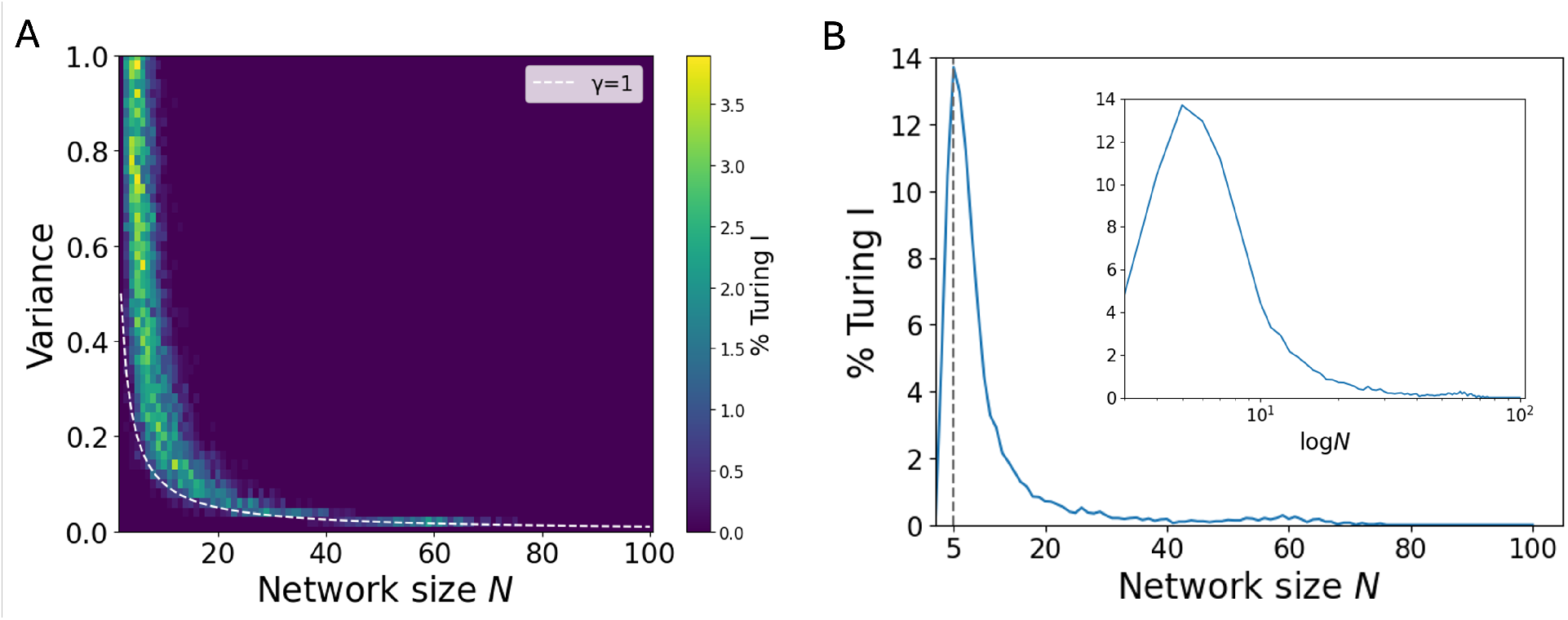
Optimal network size for highest Turing robustness. (A) Heat map of percentage occurrence of Turing I in random matrices of different networks sizes *N* and variances *σ*^2^ for *γ* = 1 and zero off-diagonal element sparsity (or connectivity *C* = 1). (B) Corresponding percentage shares of each network size *N*. For our parameters, the optimal network size is *N* = 5. The horizontal axis is set to a linear scale (logarithmic scale in inset).

Is there an optimal network size for highest robustness of the occurance of Turing patterns? We expect such an *N*_opt_ as there are no Turing patterns for *N* = 2 and for *N* → ∞. Turing patterns disappear as the eigenvalue distribution becomes a step function, allowing no instability with diffusion. To confirm our intuition, we then calculated the percentage of Turing I for each network size from the heat map (Fig. 7B), demonstrating that *N*_opt_ = 5 is the most optimal network size based on its highest percentage of Turing patterns. Thus, for a network with two diffusible species, having additional three immobile nodes gives the optimal network size for Turing instabilities. Compared to *N* = 3 with one non-diffusible species (4.73%), Turing instabilities are three times more likely to occur for *N* = 5 (13.86%).

While our random matrices are fully connected, it was suggested that biological networks tend to be sparsely connected due to its evolutionary advantage in preserving robustness [31]. Let us assume that the probability of observing *k* connections is binomially distributed 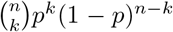 where *n* the number of matrix elements to consider (‘trials’) such as *n* = *N* (*N* − 1) for the number of off-diagonal matrix elements, and *p* the probability for having a connection (‘success’). As the variance *np*(1 − *p*) is largest for *p* = 0.5, corresponding to the maximum entropy distribution, we would expect on average 50% connections. This corresponds to the maximum number of possible network structures found by evolution. However, does sparcity also translate into higher Turing robustness? We incorporated sparsity but found that its effect is minor, without qualitatively changing our results (see Figs. S2 and S3 in Supplementary Materials). The peak value of the Turing I percentage slightly reduces with sparcity (≈ 14, 12, and 10% for respective sparcities 0, 25, and 50%), while the corresponding optimal network sizes increase slightly (*N*_opt_ = 5, 7, and 8 for above sparcities). Hence, overall the robustness of Turing is reduced with sparcity, pointing to feedback being important for pattern formation. Note the circular law also applies to sparcity, but needs modification. The radius is just rescaled to 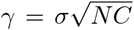 where connectivity *C* = *p* is the probability for having an off-diagonal matrix element [25]. This points to our results being broadly representative for a large range of random Turing networks.

A major constraint on Turing patterns is that the inhibitor has to diffuse much faster than the activator. How does having immobile nodes change this constraint? Previous work on small networks hinted towards a softening of this constraint [23, 22, 26]. By varying the diffusion of the two diffusible species modeled in our random reaction-diffusion systems, we sampled systems of *N* = 2, 3, 5, and 50 nodes, with *N* = 2 not supporting Turing for our random matrices with equal degradation of − 1 for all nodes (see proof in Methods). Importantly, we found that differential diffusivity is important for Turing I instabilities to arise for very few immobile nodes, but not for systems with many immobile nodes: Based on Fig. 8, a network of *N* = 3 with one immobile node has nearly zero percent Turing instabilities arising at equal diffusivity (i.e. at the identity line *D*_1_*/D*_2_ = 1). As the diffusion ratio deviates from 1, Turing instabilities become more likely for *N* = 3, creating a distinct gap around the identity line. Notably, the plot is symmetric with respect to the identity line for all *N* since the assignment to activator or inhibitor is arbitrary. In contrast, in the presence of more immobile nodes, Turing instabilities can arise at equal diffusivity: As network size increases, the gap at the identity line becomes more narrow, indicating Turing instabilities occurring at a wider combination of the diffusion parameters (see also Fig. S4 in the Supplementary Materials). For the a large network size of *N* = 50, we observed a similar occurrence of Turing instability for the majority of the diffusion parameter combinations, except for when the diffusion parameters are both extremely low, approximating the case of no diffusion. Since the Turing instability is a diffusion-driven instability, it is reasonable to have no Turing instability arising here. Additionally, there is no ‘dark line’ (indicating zero or low percentage) at the very bottom or very left of all surface plots, indicating that a Turing instability can arise in systems with only one diffusible species. The absence of differential diffusion is even stronger when consider all Turing cases (Turing I, II, and Turing-Hopf) - see Fig. S5 of the Supplementary Materials. In summary, having numerous immobile nodes significantly relaxes the constraints from Turing conditions.

**Figure 8:**
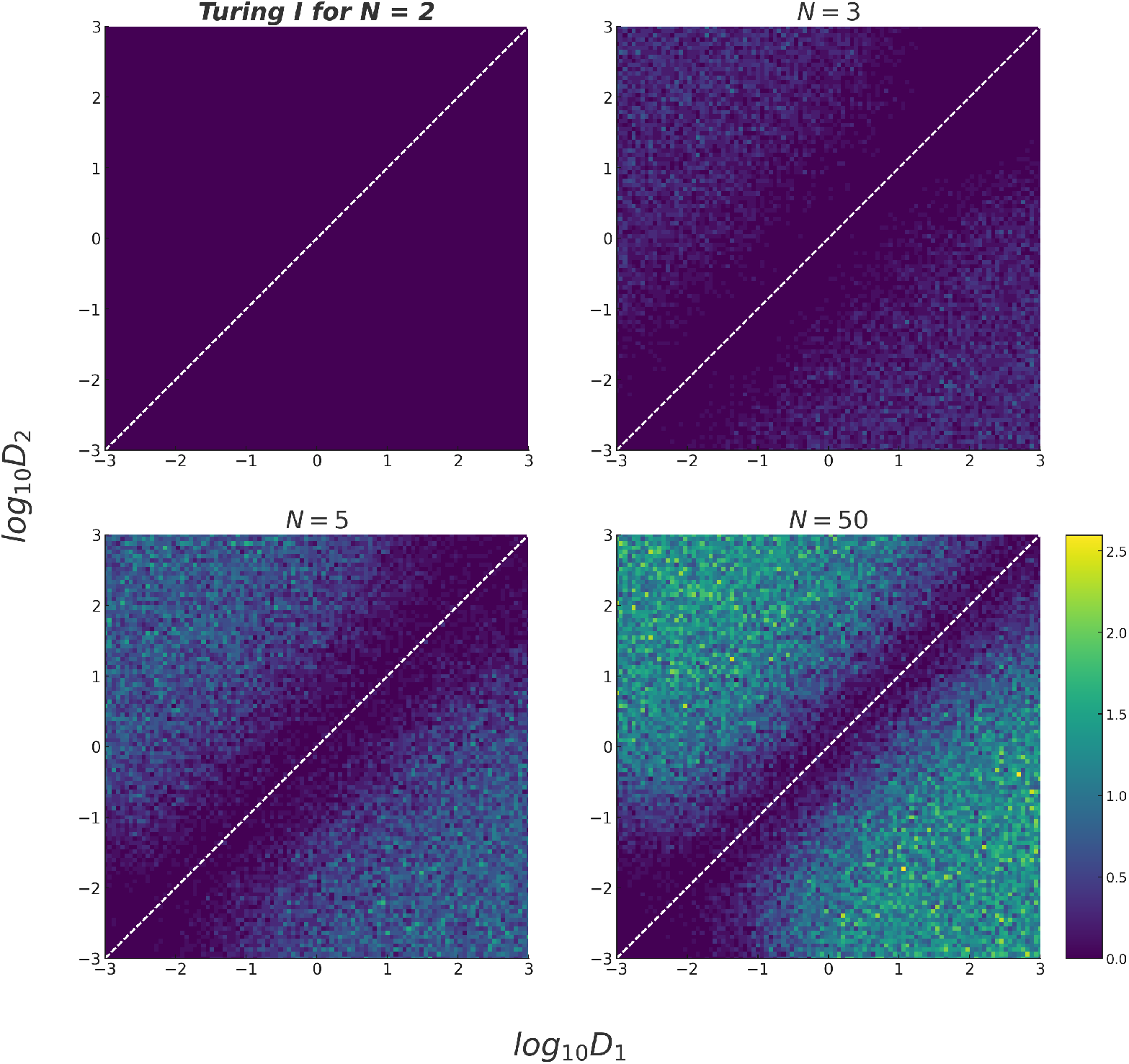
Effect of diffusion constants on Turing pattern formation. Heat map of percentage occurrence of Turing I in random matrices for different diffusion parameters, *D*_1_ and *D*_2_: (A) *N* = 2, *σ*^2^ = 0.5; (B) *N* = 3, *σ*^2^ = 0.33; (C) *N* = 5, *σ*^2^ = 0.2 (D) *N* = 50, *σ*^2^ = 0.02. For increasing *N*, constraints on diffusion constants being different vanish.

## Discussion

Turing networks have traditionally been small in size, making a direct mapping to complex developmental pathways with many unknown molecular species and parameter values difficult. Another drawback of using Turing models in biology has been the requirement to fine-tune parameters to fulfill Turing’s conditions on stability without diffusion and instability for certain wave numbers with diffusion, in contrast to biology’s robustness and evolvability [32]. Here, we circumvented these issues by using large random matrices to describe linearized Jacobian systems, greatly extending previous studies on *N* ≤ 6 [26]. By extensively sampling matrix elements, we obtained excellent statistics for networks with up to 100 molecular species.

Applying Robert May’s circular law [24] to cases without and with diffusion of two designated species, in line with Turing’s original model [4], we identified an optimal network size for maximal robustness, reaching 13.86% of sampled matrices. This value is orders of magnitude higher than previous estimates for small networks [22]. Our results are robust even when sparcity is introduced, leading only to minor changes in the observed Turing percentage or optimal network size. Importantly, an optimum in Turing robustness emerges in our model due to the tradeoff between maximizing stability without diffusion, which favors small networks centered around eigenvalues on the negative real axis, and maximizing instability with diffusion, which favors large networks for eigenvalues to reach positive real values. According to the circular law, small networks have small radii for the distribution of their eigenvalues, while large networks have large radii. For increasingly large Turing networks, we further found that differential diffusion constants are not required, in line with studies on Turing topologies [23]. Having many immobile nodes effectively allows for delays and hence different diffusion constants. We hope these results shed new light on Turing mechanisms in developmental systems and help make Turing mechanisms more relevant to quantitative biology.

To make our model interpretable, we used a number of simplifying assumptions, resulting in limitations of our predictions. A major simplification is our diagonal matrix elements, which are set to a constant value for all species, representing equal degradation rates for all species. This led to the convenient behavior that all eigenvalues are centered around a single point on the negative real axis, following Robert May’s circular law. Without that assumption, the eigenvalue distributions would still be restricted by circles, but now following the less restrictive Gerschgorin theorem. Based on the latter, the radius scales as ∼ *N*, and not 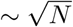 as in May’s circular law. This leads not only to larger circles but also to unions of overlapping circles, restricting the real parts less efficiently [33]. Using the same degradation rates also does not allow for flexibility in the levels of self-activation and inhibition often observed in biological pathways. Choosing degradation rates from a distribution introduces correlations among corresponding off-diagonal matrix elements *J*_*ij*_ and *J*_*ji*_ and hence different shapes of eigenvalue distributions [25].

Our work opens new avenues for exciting research in systems biology. In future work, more specific small networks could be investigated similar to the ones shown in Fig. 1, to better understand the distributions of Jacobian matrix elements with pattern-forming capability from sampled parameter values. In addition, examining random networks would allow one to obtain joint distributions of matrix elements and hence correlations among these, leading to renewed investigations into the role of network topology [28]. Of course, more diffusing species could be included, and the effects of boundary conditions investigated. The latter effect was removed in our study by considering systems of infinite domain size, in line with previous studies [22].

In conclusion, we find that Turing patterns do not only occur more frequently by chance than previously thought, but also that there is a surprising sweet spot of network size for most robust Turing patterns. This optimal network size occurred via a stability and instability tradeoff, which did not change significantly when varying the sparsity of the network. This finding supports the idea that the Turing mechanism, if employed by evolution in developmental pathways, is represented by relatively small and hence likely identifiable modules. Understanding their embedding and hierarchical organization in large developmental networks will be worthwhile endeavors.

## Methods

### Parameter searches including Latin hypercube sampling

For Fig. 3 we sampled 10^7^ parameter combinations for each network topology using Latin hypercube sampling *lhs* function from *pyDOE* library, using a loguniform distribution with the following bounds: *V*_*i*_ : (0.1, 10), *K*_*ij*_ : (0.01, 1), *b*_*i*_ : (0.001, 0.1), *µ*_*i*_ : (0.01, 1) (see Eq. 2). The diffusion constants for Species 1 and 2 were fixed to 1 and 10, and the Hill coefficient was set to *n*_*ij*_ = 2 throughout. For each parameter set we calculated the unique steady states using 10 different initial guess using the Newton-Raphson method (function *root* with ‘hybrid’ setting). In Figs. 6 and 7 we used 10^3^ random matrices for each *N* and *γ* pair, and in Fig. 8 we used 10^3^ random matrices for each *D*_1_ and *D*_2_ pair.

### Linear stability analysis

For each steady state of a parameter set, and there can be more than one due to multistability, we conducted linear stability analysis by calculating and diagonalizing the Jacobian matrices for wave numbers from 0 to *k*_*max*_ = 10 using stead size Δ*k* = 0.01 and with *linalg.eigenvals* function from *numpy* library. Subsequently, we analyzed for Turing I, II etc using a simplified version of the classification scheme in [22].

### Numerical implementation and code availability

For Fig. 3, the high-dimensional parameter space created by nested loops was flattened, and ran in parallel by dividing up the linear parameter array equally on different cores using the *multiprocessing* library. The code used for producing Figs. 2-8 can be accessed on *github.com/Endresgroup/Random Turing networks*.

## Data availability

The datasets generated and analyzed in Figs. 3 and 4 are available in the Zenodo repository https://zenodo.org/records/13142658

### Fitting of beta distribution

Fitting a beta distribution to the data of the Jacobian matrix elements in Fig. 3 involves estimating the distribution’s parameters, typically using the method of moments or maximum likelihood estimation (MLE). The method of moments involves calculating the sample mean and variance to derive initial estimates for the shape parameters *α* and *β*. MLE, on the other hand, maximizes the log-likelihood function, which measures how well the beta distribution with specific parameters explains the observed data, to find the best-fitting parameters. Both methods aim to provide a beta distribution that accurately represents the underlying data distribution. Here, we used the method of moments to obtain an initial guess for the parameter values of *α* and *β*, which we then used as an initial guess for the maximisation of the log-likelihood function.

### Mathematical proofs

Proof that the outliers of the Jacobian matrices have extreme negative real parts, following approximately 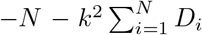. This is because the trace is given by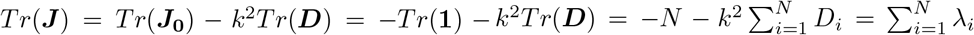, which is invariant under similarity transformations during diagonalization and hence equals the sum of all the eigenvalues *λ*_*i*_ (*i* = 1, …, *N*). Or in other words, the mean of the eigenvalues stays the same under such a transformation. Hence, at least one eigenvalue, if real, or a complex conjugate pair, if with an imaginary part, becomes an outlier and tends to minus infinity for increasing wave number *k* at a given network size *N*. This is relevant for Fig. 5A.

Proof that there are no Turing instabilities for random matrices with *N* = 2: Based on Eq. 5, the *k*-dependent Jacobian matrix is given by

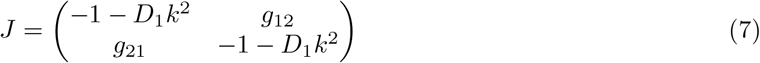

Hence, the trace is always negative for all *k*, i.e.

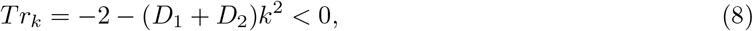

fulfilling one of Turing’s conditions. Furthermore, the determinant without diffusion (*k* = 0) is required to be positive, and hence must be

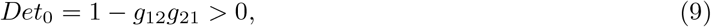

leading to the required stability in absence of diffusion. However, with diffusion (*k >* 0), the determinant cannot become negative as needed for a saddle node and hence instability. Instead, we have

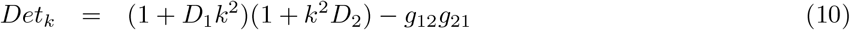

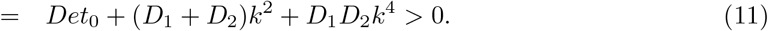

Hence, the last of Turing’s conditions cannot be fulfilled for two nodes, which is relevant for Figs. 7A and 8A.

## Acknowledgements

We thank Martina Oliver Huidobro, Mark Isalan, David Lacoste, and Roozbeh Pazuki for helpful discussions, and funding through a studentship from the Department of Life Sciences at Imperial College London (to AMG) and the AI-4-EB Consortium for Bioengineered Cells and Systems (BBSRC award BB/W013770/1) (to RGE).

