## Supplementary Material for "Optimal network sizes for most robust Turing patterns"

### Supplementary Materials

Here we show additional plots: Figue S1 extends Fig. 3A of the main text by showing empirical histograms of non-Turing Jacobian matrix elements in orange. Figures S2 and 3 extend Fig. 7 on the optimal network size for random Jacoban matrices with sparcity, i.e. missing network links, showing robustness of this key result. Figure S4 complements Fig. 8 for all Turing (instead of just Turing I), showing that there is no requirement on differential diffusion for large matrices.

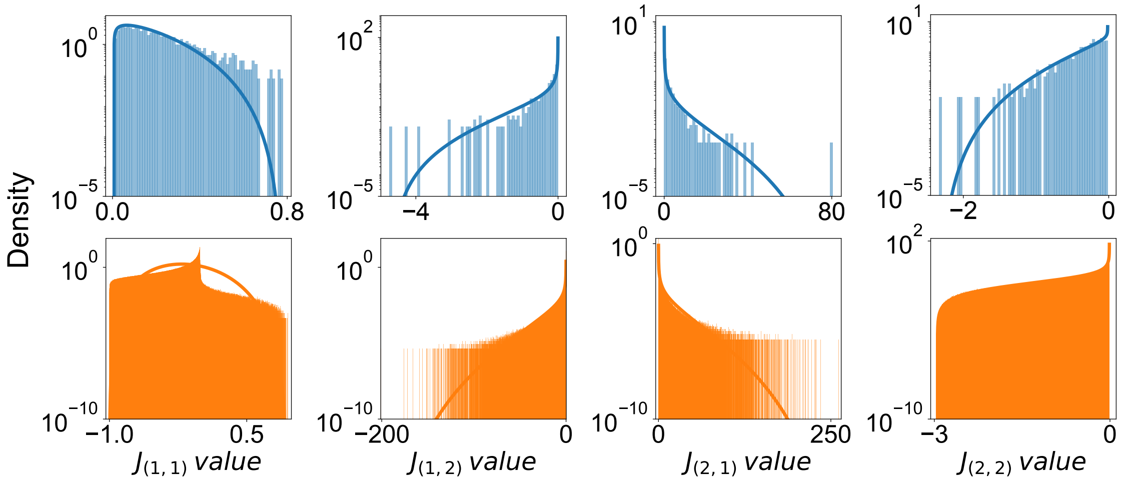


**Figure S1: Empirical distributions of Jacobian matrix elements for 2 node network.** This extends Fig. 3A, showing histograms for both Turing I instabilities (light blue) and non Turing (light orange), along with fits to beta distributions for Turing I (blue line) and non-Turing (orange line). Empirical distributions are computed based on parameter sampling for the pre-defined 2-node network topology in Fig. 1, Ai.

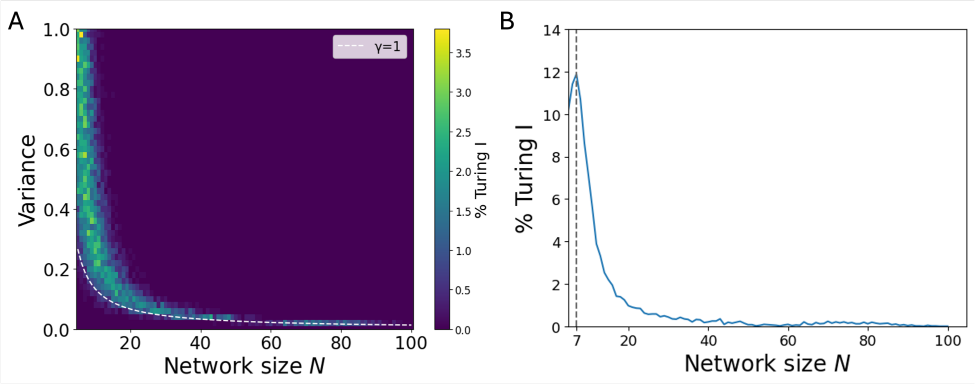


**Figure S2: Optimal network size for highest Turing robustness for small sparsity.** (A) Heat map of percentage occurrence of Turing I in random matrices of different networks size $N$ and variance $\sigma^2$ for $\gamma = 1$ and 25% off-diagonal element sparsity ($C = 0.75$). A dashed white line corresponds to the equation $\sigma^2 = 4/(3N)$. (B) Corresponding percentage shares of each network size $N$. For our parameters, the optimal network size is $N = 7$. The horizontal axis is set to linear scale.

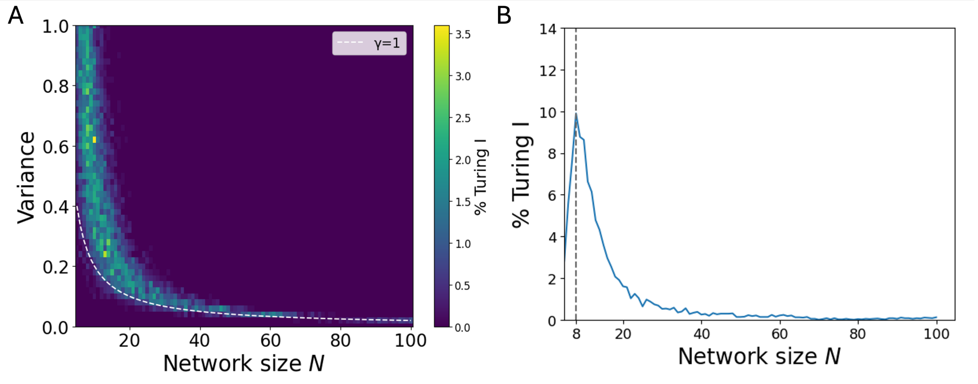


**Figure S3: Optimal network size for highest Turing robustness for strong sparsity.** (A) Heat map of percentage occurrence of Turing I in random matrices of different networks size $N$ and variance $\sigma^2$ for $\gamma = 1$ and 50% off-diagonal element sparsity ($C = 0.50$). A dashed white line corresponds to the equation $\sigma^2 = 2/N$. (B) Corresponding percentage shares of each network size $N$. For our parameters, the optimal network size is $N = 8$. The horizontal axis is set to linear scale.

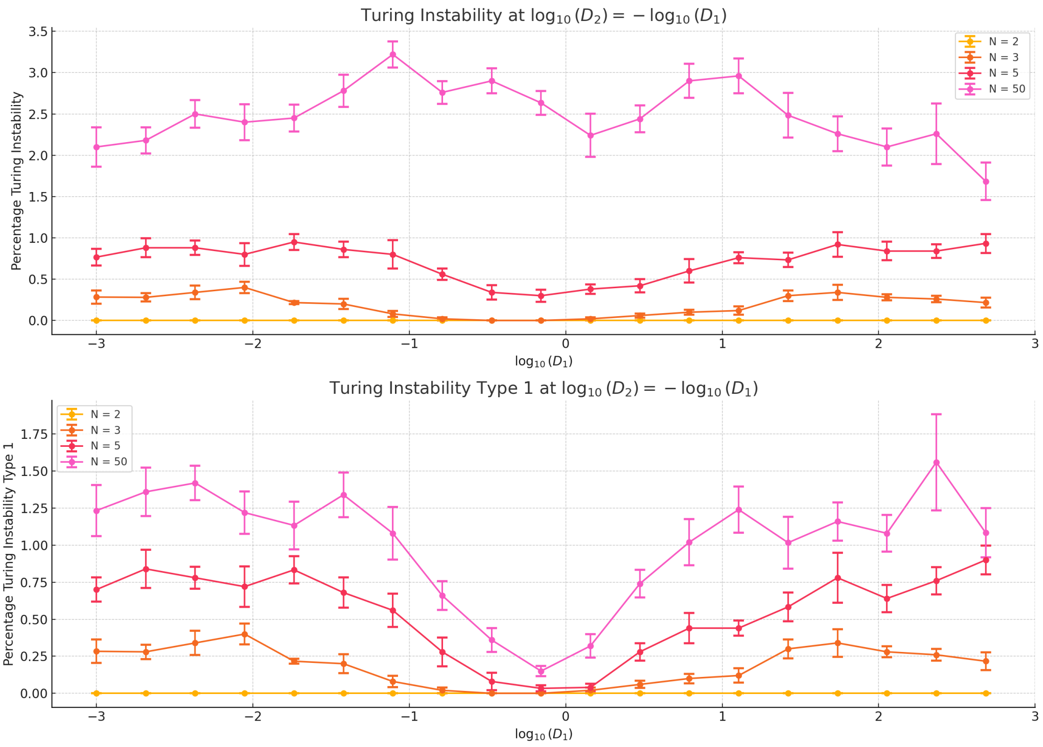


**Figure S4: Visualizing the diffusivity gap.** Profile plot for data from Figs. 8 and S5. The the heatmap simulation data was filtered and binned around the diagonal line $\log 10(D_2) = -\log 10(D_1)$, showing averages and standard errors of the Turing (top) and Turing I (bottom) percentages.

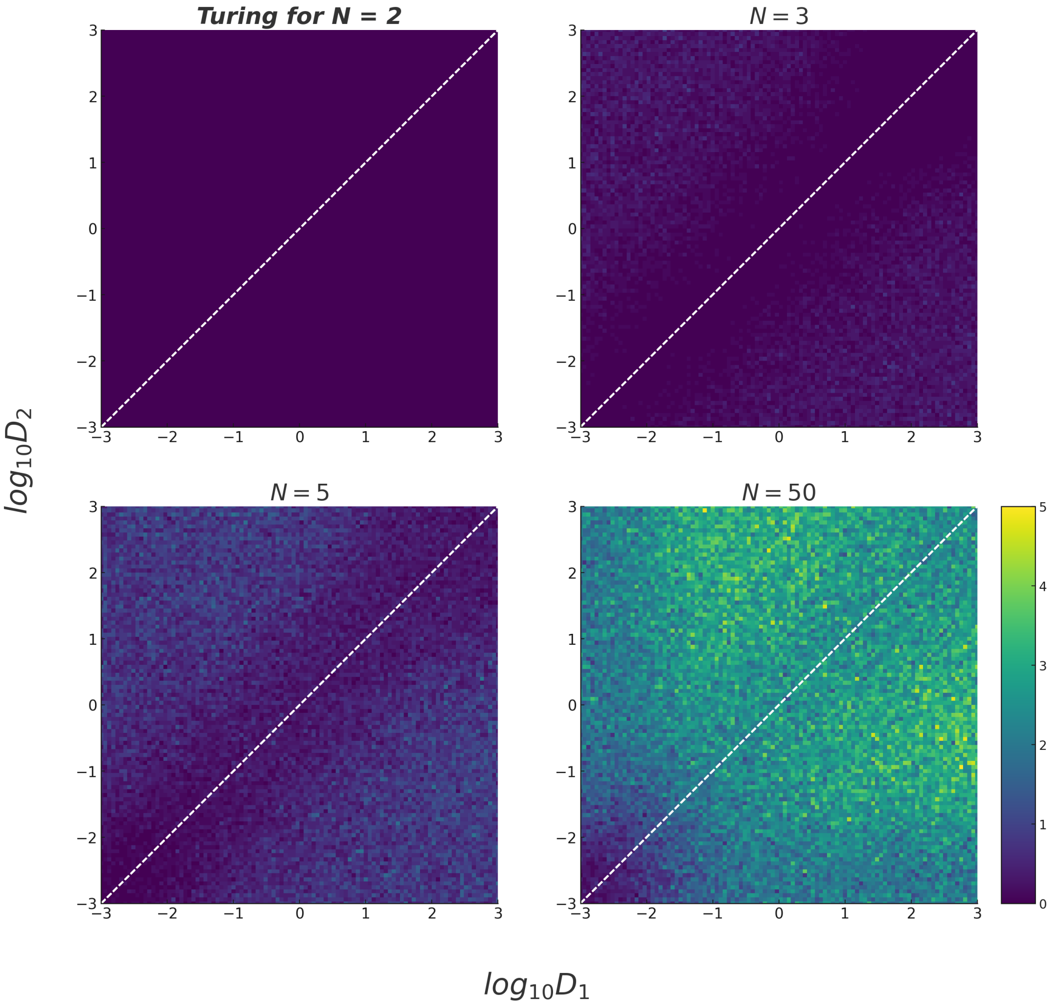


**Figure S5: Effect of diffusion constants on Turing pattern formation.** Heat map of percentage occurrence of all Turing (I, II, and Turing-Hopf) in random matrices for different diffusion parameters, $D_1$ and $D_2$: (A) $N = 2$, $\sigma^2 = 0.5$; (B) $N = 3$, $\sigma^2 = 0.33$; (C) $N = 5$, $\sigma^2 = 0.2$ (D) $N = 50$, $\sigma^2 = 0.02$. For increasing $N$, constraints on diffusion constants being different vanish.
